# fMRI-guided white matter connectivity in fluid and crystallized cognitive abilities in healthy adults

**DOI:** 10.1101/2020.01.15.907782

**Authors:** Yunglin Gazes, Jayant Sakhardande, Ashley Mensing, Qolamreza Razlighi, Ann Ohkawa, Maria Pleshkevich, Linggang Luo, Christian Habeck

## Abstract

This study examined within-subject differences among three fluid abilities that decline with age: reasoning, episodic memory and processing speed, compared with vocabulary, a crystallized ability that is maintained with age. The data were obtained from the Reference Ability Neural Network (RANN) study from which 221 participants had complete behavioral data for all 12 cognitive tasks, three per ability, along with fMRI and diffusion weighted imaging data. We used fMRI task activation to guide white matter tractography, and generated mean percent signal change in the regions associated with the processing of each ability along with diffusion tensor imaging measures, fractional anisotropy (FA) and mean diffusivity (MD), for each cognitive ability. Qualitatively brain regions associated with vocabulary were more localized and lateralized to the left hemisphere whereas the fluid abilities were associated with brain activations that were more distributed across the brain and bilaterally situated. Using continuous age, we observed smaller correlations between MD and age for white matter tracts connecting brain regions associated with the vocabulary ability than that for the fluid abilities, suggesting that vocabulary white matter tracts were better maintained with age. Furthermore, after multiple comparisons correction, the mean percent signal change for the episodic memory showed positive associations with behavioral performance, and the associations between MD and percent signal change differed by age such that, when divided into three age groups to further explore this interaction, only the oldest age group show a significant negative correlation between the two brain measures. Overall, the vocabulary ability may be better maintained with age due to the more localized brain regions involved, which places smaller reliance on long distance white matter tracts for signal transduction. These results support the hypothesis that functional activation and white matter structures underlying the vocabulary ability contribute to the ability’s greater resistance against aging.

## 1. Introduction

While aging is a pervasive process that influences most cognitive abilities, there are a number of abilities often described as crystallized, such as vocabulary and factual knowledge, that are well maintained with age past the 60’s and 70’s (Verhaegen et al., 2003). It is puzzling that the same brains can be host to both cognitive abilities that decline with age and those that remain stable. Previous studies have explored differential age-related changes in neural activities during performance of fluid abilities, such as perceptual speed and episodic memory, that were shown to decline with age (Grady et al., 2005). However, few studies have contrasted the age-related differences in brain structure and functional activation underlying fluid vs. crystallized abilities. Understanding the neural processes supporting crystallized abilities may contribute to uncovering the mechanisms that maintain neuroanatomical structures against aging.

The associations between white matter and cognitive performance have been well established in aging research. Aging studies using Diffusion Tensor Imaging (DTI), which provides a measure of white matter microstructural integrity, have shown that cognitive performance for four of the cognitive abilities is associated with DTI measures for a number of major white matter pathways (Burzynska et al., 2010; Kennedy and Raz, 2009; Madden et al., 2012; Teubner-Rhodes et al., 2016). Specifically, fractional anisotropy (FA), a DTI measure, was generally shown to be correlated positively with performance in tasks such as letter and number comparison in the prefrontal white matter (Kennedy and Raz, 2009), logical memory in the posterior internal capsule and temporal white matter (Sullivan et al., 2010) and vocabulary knowledge in the left arcuate fasciculus (Teubner-Rhodes et al., 2016). Using multivariate statistics, specific patterns of 18 major white matter tracts were shown to partially explain the effect of age on processing speed, reasoning and episodic memory (Gazes et al., 2016).

Age effects on fMRI activation across tasks are much more variable than age effects reported for white matter diffusivity. While DTI studies generally showed decreasing fractional anisotropy (FA) and increasing mean diffusivity (MD) with older age (Madden et al., 2009), functional activation studies have found both decreased and increased activations with older age across different tasks (Grady, 2008), such that the variability ranges across brain regions and tasks as well as in the direction of the associations with performance. In Cabeza et al. (2004), decreased activation was reported in the occipital cortex and increased activation in the prefrontal and parietal cortices in older than younger adults in working memory, visual attention and episodic memory tasks. The decreased activation has been attributed to worse activation capacity with older age when the processes cannot activate as much as younger adults can, and increased activation has been attributed to lower efficiency in task processing, necessitating great resources to perform the same task as younger adults (Grady, 2008; Stern, 2012).

Previous studies that combined fMRI and DTI to examine age-related changes in cognition usually focused on one or two specific cognitive tasks such as episodic (Fjell et al., 2016) and working memory (Webb et al., 2019) that tend to decline with age. However, it is essential to examine both abilities that decline (fluid abilities) as well as remain stable (crystallized abilities) with age in order to explore the mechanism underlying aging that enables certain abilities to remain stable while others decline with older age. Thus, examining four cognitive abilities that capture age-related changes are the best starting point.

While a number of brain structures and characteristics may contribute to age-related cognitive decline, the current study contrasted the brain activation and the white matter connectivity associated with vocabulary against that of three fluid abilities, episodic memory, reasoning and processing speed, as a starting point to understanding neural differences between fluid and crystallized abilities. We took advantage of data available from the Reference Ability Neural Network (RANN) study (Habeck et al., 2016), in which 12 fMRI activation tasks were administered to a large group of adults spanning 20 to 80 years old. The 12 tasks were selected from a set of cognitive tasks found to reliably estimate the four cognitive abilities examined in the study and to account for the most variance in age-related decline (Salthouse et al., 1991), and thus enabled robust approximation of brain activations underlying the four abilities. Using the brain patterns as seeds and targets, we conducted fMRI-guided tractography to extract the white matter microstructural measures for each ability.

Regionally, the vocabulary ability was expected to activate the left inferior frontal and the temporal lobe while the fluid abilities activate bilateral regions more distributed across the brain. If the functional processes and the white matter fibers supporting vocabulary are better maintained with older age than for the fluid abilities, it was hypothesized that the three brain measures, percent signal change in functional activations and FA and MD in the white matter fibers connecting the activated regions, should show significant interaction with age such that the slope of the age associations is smaller for vocabulary than for the fluid abilities.

## 2. Method

### 2.1 Participants

Healthy adult participants, n of 221, between 20-80 years old, who were strongly right-handed and native English speakers, were included in the study. Participants were recruited via random-market-mailing and screened for MRI contraindications and hearing or visual impairment that would impede testing. Older adult participants were additionally screened for dementia and mild cognitive impairment prior to participating in the study, and participants who met criteria for either were excluded. Apart from these obvious cognitive exclusion criteria, we had a host of other health-related exclusion criteria including: myocardial infarction, congestive heart failure or any other heart disease, brain disorder such as stroke, tumor, infection, epilepsy, multiple sclerosis, degenerative diseases, head injury (loss of consciousness > 5mins), mental retardation, seizure, Parkinson’s disease, Huntington’s disease, normal pressure hydrocephalus, essential/familial tremor, Down Syndrome, HIV Infection or AIDS diagnosis, learning disability/dyslexia, ADHD or ADD, uncontrolled hypertension, uncontrolled diabetes mellitus, uncontrolled thyroid or other endocrine disease, uncorrectable vision, color blindness, uncorrectable hearing and implant, pregnancy, lactating, any medication targeting central nervous system, cancer within last five years, renal insufficiency, untreated neurosyphillis, any alcohol and drug abuse within last 12 month, recent non-skin neoplastic disease or melanoma, active hepatic disease, insulin dependent diabetes, any history of psychosis or ECT, recent (past 5 years) major depressive, bipolar, or anxiety disorder, objective cognitive impairment (dementia rating scale of <130), and subjective functional impairment (BFAS > 1). Informed consent, as approved by the Internal Review Board of the College of Physicians and Surgeons of Columbia University, was obtained prior to study participation, and after the nature and risks of the study were explained.

### 2.2 fMRI activation tasks

Twelve computerized tasks, consisting of three tasks per ability, were administered to participants in two 2-hour sessions, with the order of the tasks fixed within each session, but the order of the two sessions was counterbalanced across participants.

#### 2.2.1 Episodic memory tasks

The primary dependent variable for the memory tasks was the proportion of correct trials out of all on-time trials.

##### Logical Memory

Participants were asked to answer detailed multiple-choice questions about a story presented on the computer screen, with four possible answer choices.

##### Word Order Recognition

Participants were presented with twelve words sequentially and were instructed to remember the order in which the words were presented. Following the word list, they were given a probe word at the top of the screen, and four additional word choices below. They were instructed to choose out of the four options the word that immediately followed the word given above in the word list.

##### Paired Associates

Participants were instructed to remember pairs of words presented sequentially on the screen. Following presentation of the pairs, participants were given a probe word at the top of the screen and four additional word choices below. Participants were asked to choose the word that was originally paired with the probe word.

#### 2.2.2 Reasoning tasks

The primary dependent variable for the reasoning tasks was proportion of correct trials out of all on-time trials.

##### Paper Folding (Ekstrom et al., 1976)

Participants selected which of 5 options best represented the pattern of holes that would result from a sequence of folds in a piece of paper through which a hole was punched. The sequence was given on the top of the screen, and the six options were given across two rows (three options in each row) below. Response consisted of pressing 1 of 5 buttons corresponding to the chosen solution.

##### Matrix Reasoning (adapted from (Raven, 1962))

Participants were given a matrix that was divided into nine cells, in which the figure in the bottom right cell was missing. Participants were instructed to evaluate which of eight figure choices, presented below the matrix, would best complete the missing cell.

##### Letter Sets (Ekstrom et al., 1976)

Participants were presented with five sets of letters, where four out of the five sets had a common rule (i.e. they have no vowels), with one of the sets not following this rule. Participants identified the unique set.

#### 2.2.3 Processing speed Tasks

As accuracy for all three speed tasks was high, the primary dependent variable was reaction time (RT) for on-time correct responses. For all tasks, participants were instructed to respond as quickly and accurately as possible.

##### Digit Symbol

A code table was presented on the top of the screen, consisting of 9 number (ranging in value from 1-9)-symbol pairs. Below the code table, an individual number/symbol pair was presented. Participants indicated whether the individual pair was the same as that in the code table.

##### Letter Comparison (Salthouse et al., 1991)

Two strings of letters, each consisting of three or six letters, were presented alongside one another. Participants indicated whether the letter-strings were the same or different.

##### Pattern Comparison (Salthouse et al., 1991)

Two figures, consisting of varying numbers of lines connecting at different angles, were presented alongside one another. Participants indicated whether the figures were the same or different.

#### 2.2.4 Vocabulary Tasks

The primary dependent for all vocabulary tasks was the proportion of correct responses.

##### Synonyms (Salthouse, 1993)

Participants were instructed to match a given probe word to its synonym or to the word most similar in meaning. The probe word was presented in all capital letters at the top of the screen, and four numbered choices were presented below. Participants indicated which choice was correct.

##### Antonyms (Salthouse, 1993)

Participants matched a given word to its antonym, or to the word most different in meaning. The probe word was presented in all capital letters at the top of the screen, and four numbered choices were presented below. Participants indicated which choice was correct.

##### Picture Naming (Woodcock et al., 1989)

Participants verbally named pictures, adapted from the picture naming task of the WJ-R Psycho-Educational battery.

### 2.3 Image acquisition procedures

All MR images were acquired on a 3.0 Tesla Philips Achieva Magnet. There were two, 2-hour MR imaging sessions to accommodate the 12 fMRI activation tasks as well as the additional imaging modalities, described below. At each session, a scout, T1-weighted image was acquired to determine participant position. Participants underwent a T1-weighted MPRAGE scan to determine brain structure, with a TE/TR of 3/6.5 ms and Flip Angle of 8 degrees, in-plane resolution of 256 ×256, field of view of 25.4 × 25.4 cm, and 165~180 slices in axial direction with slice-thickness/gap of 1/0 mm. All scans used a 240 mm field of view. For the EPI acquisition (BOLD fMRI), the parameters were: TE/TR 20/2000 ms, Flip Angle 72°, In-plane resolution 112×112 voxels, Slice thickness/gap 3/0 mm and 41 slices. Diffusion weighted images (DWI) were acquired in 55 directions using these parameters: b = 800 s/mm2, TE = 69 ms, TR = 11032 ms, Flip Angle = 90°, in-plane resolution 112 × 112 voxels, 2×2×2mm voxels, 75 slices, acquisition time 12 min 56 sec. In addition, FLAIR, ASL and a 7-minute resting BOLD scan were acquired. A neuroradiologist reviewed each participant’s scans. Any significant findings were conveyed to the participant’s primary care physician.

### 2.4 fMRI preprocessing

Each participant’s 12 task-activation fMRI scans were pre-processed in FSL (Smith et al., 2004) using the following steps: (1) within-participant histogram computation for each participant volume to identify noise (FEAT)(Woolrich et al., 2001); (2) participant-motion correction (MCFLIRT)(Jenkinson et al., 2002); (3) slice-timing correction; (4) brain-mask creation from first volume in participant’s fMRI data; (5) high-pass filtering (T=128 sec); (6) pre-whitening; (7) General-Linear-Model (GLM) estimation with equally temporally filtered regressors and double-gamma hemodynamic response functions; and (8) registration of functional and structural images with subsequent normalization into MNI space (FNIRT)(Andersson et al., 2010). General linear models (GLM) for each participant and each task consisted of block-based time-series analysis for speed and vocabulary tasks, and event-related modeling for reasoning and memory tasks. For memory tasks, only the recognition phase of the trial was analyzed. For group analysis, the block contrast was used for all of the tasks except for memory tasks, in which only the recognition contrast was used.

### 2.5 Ability-unique brain activations

For each participant, to generate a Fisher’s z-map for each ability, the twelve task contrasts were correlated with a 12-element vector selective for the particular ability. For example, for reasoning z-map, the vector would have 1’s for the three reasoning tasks and 0’s for all other tasks. This will generate a Fisher’s Z-map from transforming r to Z values at each voxel, in which the voxels with the highest correlation value for reasoning tasks have the highest Z values. At the group level, t-tests were performed for each ability across all participants, resulting in t-maps showing voxels most selective for each ability. To correct for multiple comparisons, False Discovery Rate (Benjamini and Hochberg, 1995) was applied to obtained q values of .01, a value low enough to ensure robust results. The thresholded maps were then further thresholded with a cluster size of 500 voxels to isolate clusters large enough to enable unique white matter tracts to be extracted. To create the ability-unique maps, voxels present in the overlap between any pairing of the ability maps were eliminated from each ability map, resulting in four maps in which the clusters are uniquely associated with each ability. Means for percent signal change were extracted from these ability-unique ROIs using Featquery (Woolrich et al., 2004) and examined in subsequent statistical tests.

### 2.6 Diffusion Weighted Imaging

Diffusion weighted imaging data were processed in MRtrix3 (www.mrtrix.org) using MRtrix tools except where indicated. Preprocessing was done in the following order: (1) denoising (Veraart et al., 2016), (2) Gibbs ring correction (Kellner et al., 2016), (3) corrections for motion and eddy currents (FSL eddy) (Andersson and Sotiropoulos, 2016) and (4) bias field correction (Tustison et al., 2010). Data were then modeled with Constrained Spherical Deconvolution (Tournier et al., 2004) from which the fiber orientation distribution were estimated for white matter and the cerebrospinal fluid (CSF) using the multi-shell multi-tissue approach adopted for a single non-zero b-value along with a b-zero volume. Then probabilistic tractography was performed using tckgen function in MRtrix3 (Tournier et al., 2010) for the whole brain in each participant, using FSL’s grey/white matter and CSF segmented images as anatomical constraints, to generate 20 million fiber tracts with a maximum angle of 45 degrees and maximum length of 250mm. Streamlines were corrected for bias with the SIFT2 algorithm (Smith et al., 2015) in MRtrix3. Then, to extract the white matter tracts connecting each pair of ROIs in the ability-unique activation maps, tckedit function was used to select streamlines that connect pairs of ROIs. The resulting tracts were then used as masks to extract FA and MD for each ability in each participant. Due to the inexact nature of probabilistic tractography, some streamlines that stray outside of the main paths are unavoidable unless manual adjustment of individualized tractography parameters are used, which is not feasible for hundreds of participants and may introduce bias due to unequal treatment across participants. To address this, median FA/MD was used to extract a representative FA/MD value for the streamlines connecting each region pair. Then for each ability, the means of the FA/MD values were calculated across the white matter tracts connecting the region pairs.

Only FA and MD were examined in our study and did not include axial and radial diffusivity values because factor analysis showed that the four DTI measures are so highly correlated that they constitute at most two unique factors for each of the 18 major white matter tracts examined (unpublished result). Therefore, we only examined two measures that dominated the two factors to minimize the chance of type I and type II errors in hypothesis testing: (1) If we didn’t correct for multiple comparisons and thus overestimating the effects, then we would risk producing false positives, and (2) if we corrected for multiple comparisons, which assumes the measures are independent of each other, then we would risk producing false negatives. Therefore, we examined two measures that are more representative of the two major sources of variability in the four DTI measures and corrected for multiple comparisons as described in the Statistical analysis section below.

### 2.7 Statistical analyses

In order to compare the three brain measures, percent signal change, FA and MD, across the four abilities, which may be different purely due to natural variability across brain regions, each of the measures was converted to z-scores for each ability across the entire sample. All analyses were done in SPSS (IBM v26) unless indicated otherwise and p<.05 two-tailed corrected for multiple comparisons using Bonferroni correction.

#### 2.7.1 Ability differences in associations with age

To test if the age associations for the three brain measures differ across the four abilities, repeated-measures ANOVAs were performed for each pairing of the four abilities, in which continuous age was a covariate and two abilities were the repeated-measures factor, and the Age by Ability interaction was the effect of interest. Pairs of the abilities were tested in each model in order to examine which pairings are significantly different in their associations with age, resulting in six different models tested for each brain measure. Any Age by Ability interaction with p < 0.00278 (3 measures × 6 possible pairings) was considered statistically significant.

#### 2.7.4 Brain activation and white matter measures as explanatory variables for behavior

To examine if percent signal change in brain activation and the FA and MD for the associated white matter tracts explain the variability in the mean z-scores for the behavioral performance in each cognitive ability, the mean percent signal change and one of the white matter measures (FA/MD) along with continuous age, sex, and education were entered as explanatory variables and the respective mean z-score was entered as the dependent variable in each model. Two interaction terms, age by percent signal change and age by white matter, were also entered into each model. Any non-significant interactions were dropped out to form a reduced model, which was the final model that was run. Percent signal change, FA and MD were all converted to z-scores before entering into each statistical model.

#### 2.7.5 White matter measures as explanatory variables for brain activation

To examine if white matter measures contribute to the variability in the percent signal change in brain activation for each ability, the mean white matter measure (FA/MD), continuous age, sex, education and age by white matter measure were the explanatory variables with the mean percent signal change as the dependent variable. If the interaction term was not significant, then the reduced model was tested. Percent signal change, FA and MD were all converted to z-scores before entering into each statistical model.

## 3. Results

### 3.1 Participants

Of the 221 participants, two were missing vocabulary task scores due to software error and one had an outlier value for vocabulary MD. Table 1 shows the demographics of the full participant set divided into tertiles (Group 1: age<47 years; Group 2: 47< age < 64.74years; Group 3: age > 64.74 years). Sample sizes were not perfectly equal due to multiple participants having the same age at the tertiles. Differences across the three age groups was only significant for Age as expected, F(2, 220) = 781.6, p<.001, and not for the other three demographic variables (Education: F(2,220) = 2.00, p = .138; Sex: F(2, 220) = .045, p = .956; and DRS: F (2, 220) = .956, p = .386). Continuous age was used for most of the analyses; age groups were used for examining interaction effects with age.

**Table 1.** Participant demographic information.

| Age Group | N | Age (SD) | Sex (F/M) | Edu (SD) | DRS total |
| --- | --- | --- | --- | --- | --- |
| 1 | 72 | 34.7 (.834) | 40/32 | 16.1 (.321) | 139.8 (.327) |
| 2 | 76 | 56.5 (.611) | 42/34 | 16.0 (.250) | 140.3 (.313) |
| 3 | 73 | 71.4 (.486) | 42/31 | 16.8 (.301) | 139.7 (.327) |

### 3.2 Activation patterns for the four abilities

Each of the ability activation patterns consisted of distinct brain regions consistent with previous studies. Figure 1 displays the brain regions unique to each cognitive ability in different colors. As expected, reasoning recruited the most dorsal and lateral frontal regions, including bilateral superior and middle frontal gyri, and extending to the medial frontal gyrus and posteriorly to large portions of the precuneus, bilateral supramarginal gyri down to the superior and middle temporal gyri and small area in the right hippocampus (MNI 28, −28, −10). Memory regions included the left parahippocampal gyrus (MNI −27, −38, −10) down to the Culmen of the Cerebellum, left middle to superior temporal gyri extending anteriorly to the temporal pole, bilateral anterior cingulate extending anteriorly to the medial frontal gyrus, bilateral precentral gyri, the right postcentral gyrus and parts of the thalamus. Speed was associated with large subcortical regions such as the bilateral caudate, nucleus accumbens, thalamus and brain stem, and also included bilateral anterior cingulate, bilateral superior portions of the lateral occipital cortices, the cuneus, the right precentral gyrus extending anteriorly to the middle frontal gyrus, bilateral orbital frontal cortices and the insular cortices. Vocabulary regions were much more localized to the left hemisphere and consisted of large portions of the left inferior frontal gyrus, the precentral gyrus, the superior and middle temporal gyri and bilateral cerebellum. Overall, these regions were consistent with the functions attributed to each brain region according to the existing literature.

**Figure 1.**
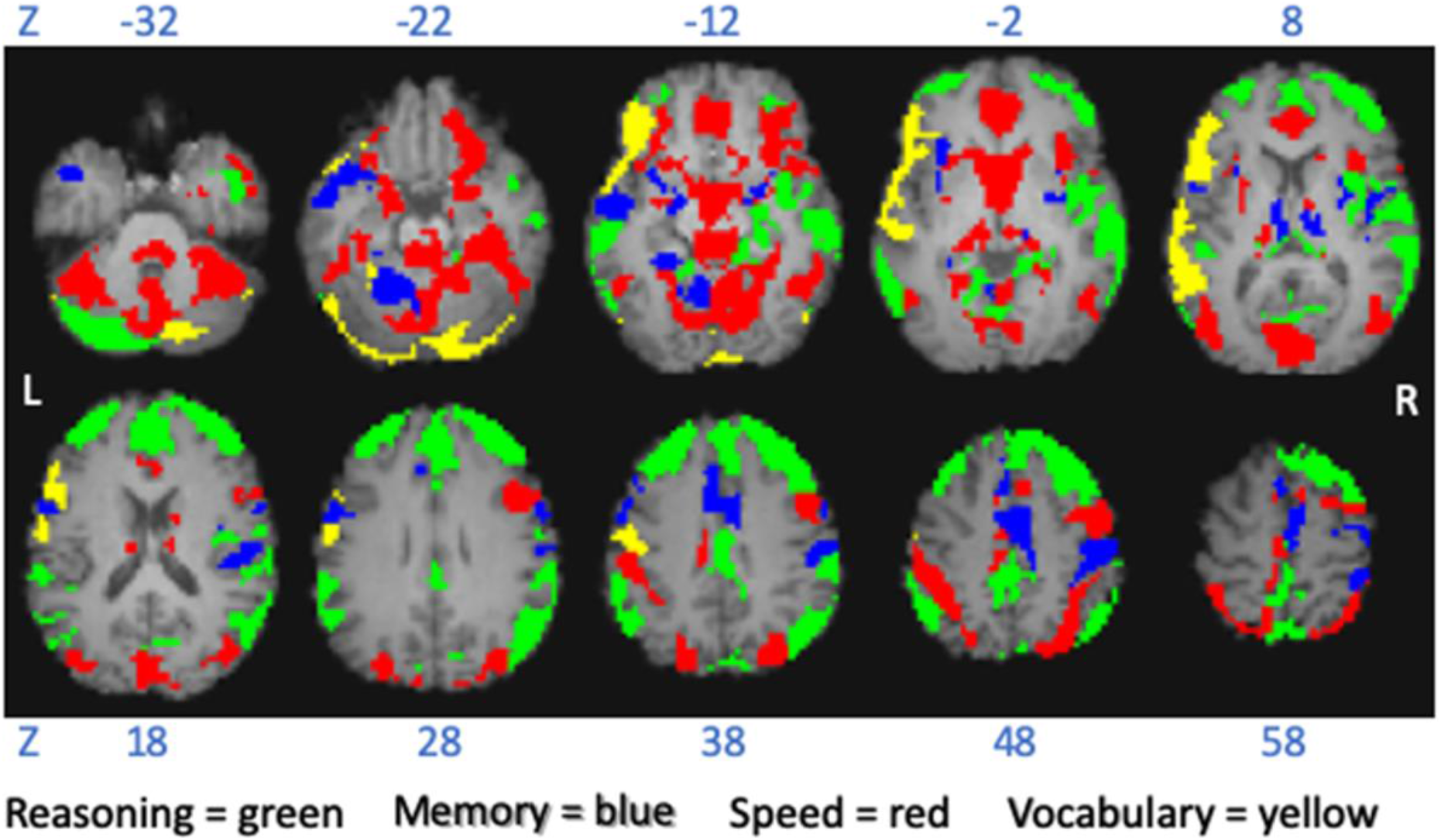
Activation patterns for the four cognitive abilities examined in the study. Z = MNI coordinate; L = left; R = right

### 3.3 White matter tracts connecting the ability-unique regions

Figure 2 shows two examples of the tracts obtained from tractography for a randomly selected participant. As can be seen in the figure, while the majority of the streamlines were as expected based on each individual’s anatomical structure, especially for the speed tracts, there were streamlines that strayed from the main path. As already stated in the Method section, this is one of the known disadvantages of tractography that would require extensive supervision at the individual level to eliminate. To minimize the influence of these extraneous streamlines on the mean FA/MD for each white matter tract, median value was used instead of the mean for the FA/MD across the streamlines in each white matter tract. Summarizing across white matter pathways connecting all of the brain regions for each ability was still performed with the mean.

**Figure 2.**
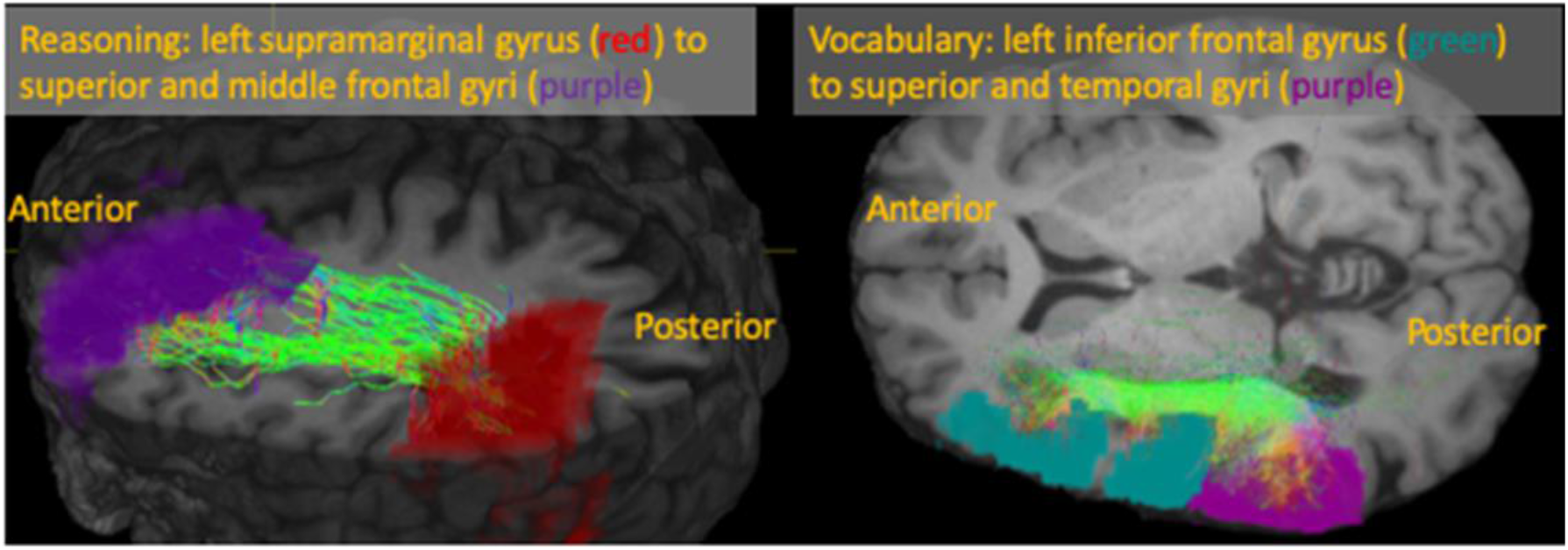
Examples of white matter tracts connecting region pairs for reasoning and vocabulary. The colors show the direction of the streamlines

### 3.4 Ability differences in correlations with age

Correlations between age and each of the behavioral and brain measure z-scores are listed in Table 2 and plotted in Figure 3. As expected, with older age, behavioral z-scores showed decreasing accuracy for Memory and Reasoning and longer RT for Speed. Also as expected, Vocabulary was the only ability that showed increasing accuracy with older age. Percent signal change in the ability-unique activated regions for each ability showed positive correlations with age for reasoning, speed and vocabulary, but memory did not show any significant association with age. White matter measures showed the expected age trend of worsening diffusivity with older age for both FA and MD for all four abilities examined, but mean FA for memory did not reach significance after correcting for multiple comparisons.

**Table 2.** Pearson r-values between each behavioral/brain measure and age for the four abilities.

|  | Fluid | Memory | Speed | Vocabulary |
| --- | --- | --- | --- | --- |
| Behavior | **-0.254\*\*\*** | **-.206\*\*** | **.509\*\*\*** | **.367\*\*\*** |
| %SigChg | **.209\*\*** | -.037 | **.345\*\*\*** | **.447\*\*\*** |
| FA | **-.402\*\*\*** | -.147* | **-.369\*\*\*** | **-.248\*\*\*** |
| MD | **.447\*\*\*** | **.432\*\*\*** | **.433\*\*\*** | **.281\*\*\*** |

Note. %SigChg=percent signal change. ^p<.08, \*p<.05, \*\*p<.01, \*\*\*p<.001, bolded: p<.003125, corrected for multiple comparison.

**Figure 3.**
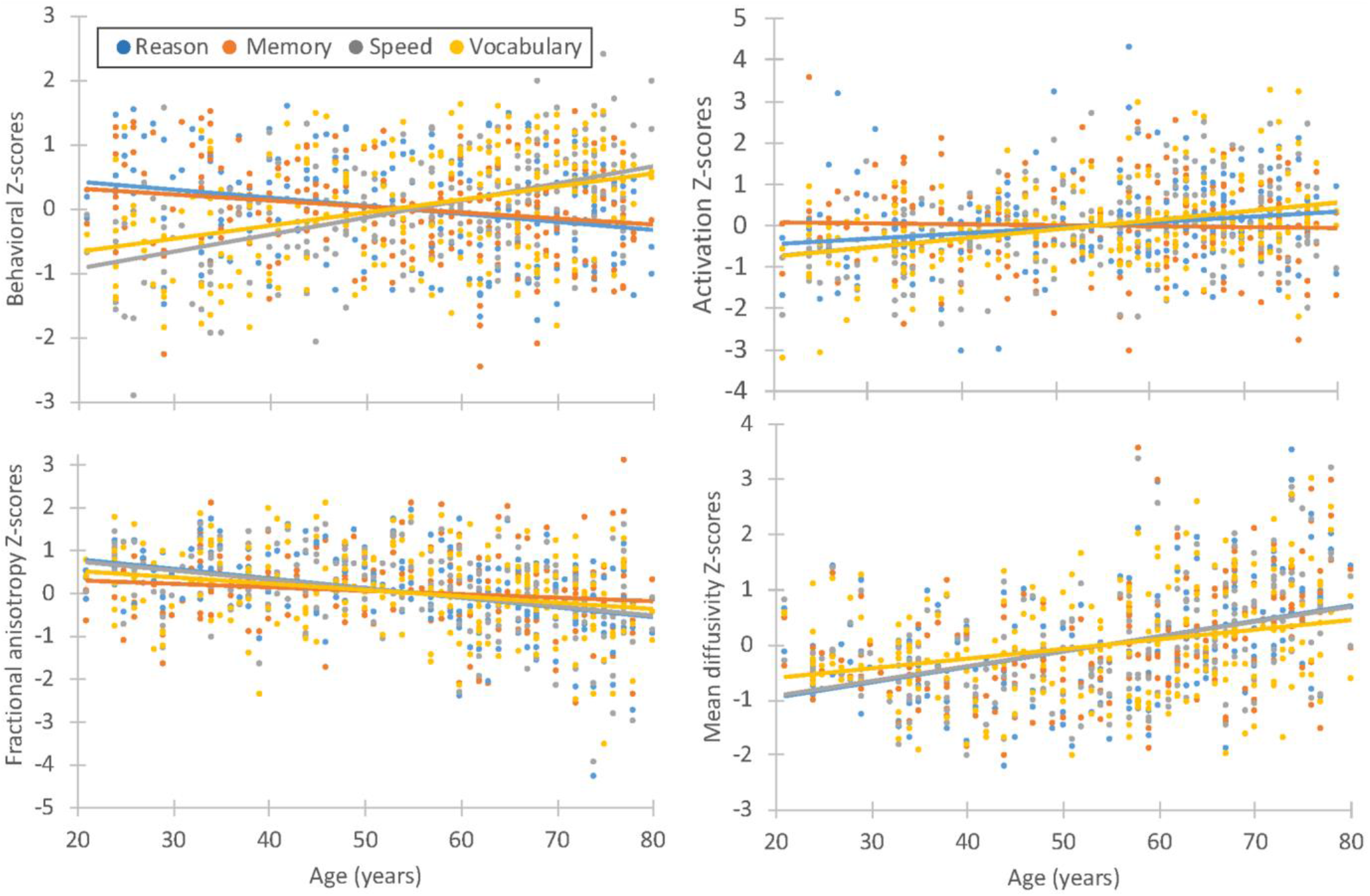
Scatter plots of z-scores for behavioral and brain measures against increasing age. Note the differences in scale across plots. Behavioral z-scores for speed measure reaction time and thus reaction time increases with older age. Behavioral z-scores for reasoning, memory and vocabulary measure accuracy. Vocabulary was the only ability that showed better performance with older age. For activation, memory region did not show any associations with age while brain regions for the other abilities increased in activation with older age. Fractional anisotropy for vocabulary-related white matter tracts were similar in age associations as the fluid abilities. Mean diffusivity for vocabulary-related white matter tracts showed the smallest correlation with age while the fluid abilities exhibited very similar age correlations such that the three linear lines are lined up on top of each other.

To test if the associations with age differs across abilities, age by ability interaction was examined for each of the three brain measures, percent signal change, FA and MD, for each of the six possible parings among the four abilities. Table 3 listed the F values for the Ability variable and the Age by Ability interaction. The percent signal change for memory showed an age trend that was significantly different from all other abilities such that it does not show an association with age while vocabulary’s percent signal change showed the highest correlation with age. FA for memory also showed distinct age correlations from reasoning and speed but not from vocabulary since it showed the weakest negative correlation with age, followed by the stronger association for vocabulary. For MD, correlations with age for vocabulary brain measures were the most consistent with the hypothesized direction in that vocabulary showed the weakest positive correlation with age relative to the other three abilities.

**Table 3.** F-values for Ability and Age by Ability for each brain measure for the six pairings of the four abilities.

|  | Ability |  |  | Age x Ability |  |  |
| --- | --- | --- | --- | --- | --- | --- |
|  | %SigChg | FA | MD | %SigChg | FA | MD |
| *Vs. Vocabulary* |  |  |  |  |  |  |
| **Memory** | <.001 | <.001 | 0.031 | **25.7\*\*\*** | 2.35 | **14.3\*\*\*** |
| **Reasoning** | <.001 | <.001 | 0.0088 | 3.01 | 6.63* | **18.8\*\*\*** |
| **Speed** | <.001 | <.001 | 0.0892 | 0.006 | 4.40* | **16.4\*\*\*** |
| *Vs. Memory* |  |  |  |  |  |  |
| **Reasoning** | <.001 | <.001 | <.001 | 7.95** | **40.9\*\*\*** | 0.361 |
| **Speed** | <.001 | <.001 | <.001 | **21.6\*\*\*** | **28.4\*\*\*** | 0.002 |
| *Vs. Reasoning* |  |  |  |  |  |  |
| **Speed** | <.001 | <.001 | <.001 | 3.16^ | 1.19 | 0.494 |

Note. %SigChg=percent signal change, FA=fractional anisotropy, MD=mean diffusivity. ^ $p < .08$ , \* $p < .05$ , \*\* $p < .01$ , \*\*\* $p < .001$ , bolded: $p < .00278$ , corrected for multiple comparison.

### 3.5 Brain activation and white matter measures as explanatory variables for behavior

Statistics for the 8 models (4 abilities × 2 white matter measures) are shown in Table 4. None of the interactions with age group was significant. In the reduced model, percent signal change was significant for memory with both FA and MD as the white matter measures, showing that percent signal change explains some of the variability in memory performance. Percent signal change and white matter measures did not explain behavioral variability for reasoning, speed and vocabulary.

**Table 4.**
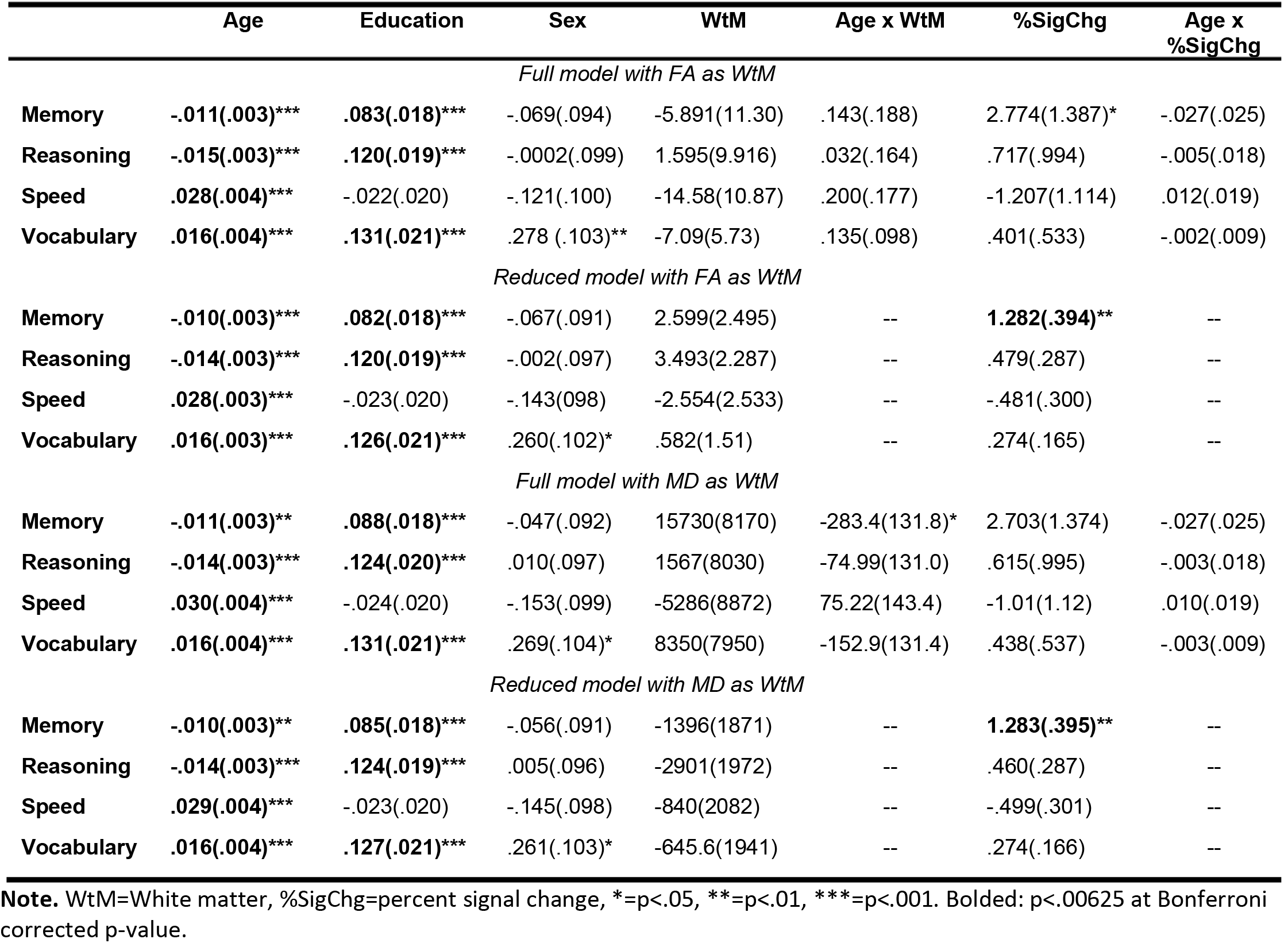
Parameter estimates (standard errors) for models with independent variables across the top and each respective ability’s behavioral score as the dependent variable. Top two sections show FA as the white matter measure and bottom two sections show MD as the white matter measure.

|  | Age | Education | Sex | WtM | Age x WtM | %SigChg | Age x %SigChg |
| --- | --- | --- | --- | --- | --- | --- | --- |
| *Full model with FA as WtM* |  |  |  |  |  |  |  |
| Memory | -.011(.003)*** | .083(.018)*** | -.069(.094) | -5.891(11.30) | .143(.188) | 2.774(1.387)* | -.027(.025) |
| Reasoning | -.015(.003)*** | .120(.019)*** | -.0002(.099) | 1.595(9.916) | .032(.164) | .717(.994) | -.005(.018) |
| Speed | .028(.004)*** | -.022(.020) | -.121(.100) | -14.58(10.87) | .200(.177) | -1.207(1.114) | .012(.019) |
| Vocabulary | .016(.004)*** | .131(.021)*** | .278 (.103)** | -7.09(5.73) | .135(.098) | .401(.533) | -.002(.009) |
| *Reduced model with FA as WtM* |  |  |  |  |  |  |  |
| Memory | -.010(.003)*** | .082(.018)*** | -.067(.091) | 2.599(2.495) | -- | **1.282(.394)\*\*** | -- |
| Reasoning | -.014(.003)*** | .120(.019)*** | -.002(.097) | 3.493(2.287) | -- | .479(.287) | -- |
| Speed | .028(.003)*** | -.023(.020) | -.143(.098) | -2.554(2.533) | -- | -.481(.300) | -- |
| Vocabulary | .016(.003)*** | .126(.021)*** | .260(.102)* | .582(1.51) | -- | .274(.165) | -- |
| *Full model with MD as WtM* |  |  |  |  |  |  |  |
| Memory | -.011(.003)** | .088(.018)*** | -.047(.092) | 15730(8170) | -283.4(131.8)* | 2.703(1.374) | -.027(.025) |
| Reasoning | -.014(.003)*** | .124(.020)*** | .010(.097) | 1567(8030) | -74.99(131.0) | .615(.995) | -.003(.018) |
| Speed | .030(.004)*** | -.024(.020) | -.153(.099) | -5286(8872) | 75.22(143.4) | -1.01(1.12) | .010(.019) |
| Vocabulary | .016(.004)*** | .131(.021)*** | .269(.104)* | 8350(7950) | -152.9(131.4) | .438(.537) | -.003(.009) |
| *Reduced model with MD as WtM* |  |  |  |  |  |  |  |
| Memory | -.010(.003)** | .085(.018)*** | -.056(.091) | -1396(1871) | -- | **1.283(.395)\*\*** | -- |
| Reasoning | -.014(.003)*** | .124(.019)*** | .005(.096) | -2901(1972) | -- | .460(.287) | -- |
| Speed | .029(.004)*** | -.023(.020) | -.145(.098) | -840(2082) | -- | -.499(.301) | -- |
| Vocabulary | .016(.004)*** | .127(.021)*** | .261(.103)* | -645.6(1941) | -- | .274(.166) | -- |

**Note.** WtM=White matter, %SigChg=percent signal change, \*=p<.05, \*\*=p<.01, \*\*\*=p<.001. Bolded: p<.00625 at Bonferroni corrected p-value.

### 3.6 White matter measures as explanatory variables for brain activation

Only white matter tracts associated with speed showed significant interactions between age and MD in explaining variability in percent signal change. Table 5 lists the parameter estimates for the variables in each group. To explore this interaction, percent signal change was plotted against MD for each of the age groups in Figure 4. The oldest age group’s percent signal change for speed regions showed a negative association with MD (r = −.310, p = .008), demonstrating that higher brain activation was associated with better white matter integrity, for speed tracts, while the two younger groups did not show significant associations between percent signal change and MD (young group: r = .128, middle age group: r = .124, both p > .05).

**Table 5.** Parameter estimates (standard errors) for models with independent variables across the top and each respective percent signal change z-score as dependent variable.

|  | Age | Education | Sex | WtM | Age x WtM |
| --- | --- | --- | --- | --- | --- |
| *Full model with FA as WtM* |  |  |  |  |  |
| Memory | -.00360(.00428) | .0605(.0272)* | -.231(.140) | .0192(.0721) | -.00456(.00455) |
| Reasoning | **.0148(.00454)\*\*** | -.0322(.0265) | .00218(.136) | .0521(.0758) | .00181(.00452) |
| Speed | **.0227(.00430)\*\*\*** | -.0181(.0256) | -.103(.131) | .0281(.0728) | .00797(.00428)^ |
| Vocabulary | **.0221(.00407)\*\*\*** | .0437(.0257) | -.0974(.128) | -.0833(.0655) | .00252(.00401) |
| *Reduced model with FA as WtM* |  |  |  |  |  |
| Memory | -.00371(.00428) | .0611(.0272)* | -.201(.137) | -.00297(.0686) | -- |
| Reasoning | **.0151(.00448)\*\*** | -.0331(.0264) | -.0126(.133) | .0614(.0721) | -- |
| Speed | **.0237(.00430)\*\*\*** | -.0221(.026) | -.181(.128) | .0762(.0684) | -- |
| Vocabulary | **.0222(.00407)\*\*\*** | .0410(.0253) | -.109(.127) | .0891(.0647) | -- |
| *Full model with MD as WtM* |  |  |  |  |  |
| Memory | -.00526(.00468) | .0598(.0273)* | -.189(.137) | .0696(.0814) | -.00260(.00532) |
| Reasoning | **.0134(.00464)\*\*** | -.0343(.0271) | -.0416(.135) | .0289(.789) | -.00291(.00494) |
| Speed | **.0230(.00432)\*\*\*** | -.0151(.0253) | -.101(.127) | .0135(.0748) | **-.0149(.00466)\*\*** |
| Vocabulary | **.0224(.00415)\*\*\*** | .0477(.0260)^ | -.0626(.129) | -.0712(.0697) | -.00126(.00454) |
| *Reduced model with MD as WtM* |  |  |  |  |  |
| Memory | -.00503(.00465) | .0589(.0272)* | -.198(.135) | .0540(.0747) | -- |
| Reasoning | **.0134(.00464)\*\*** | -.0350(.0270) | .0345(.134) | .0141(.0747) | -- |
| Speed | **.0240(.00440)\*\*\*** | -.0182(.0258) | -.157(.128) | -.0760(.0708) | -- |
| Vocabulary | **.0224(.00414)\*\*\*** | .0470(.0258)^ | -.0659(.128) | -.0766(.0668) | -- |

Note. WtM=White matter, ^p<.08, \*p<.05, \*\*p<.01, \*\*\*p<.001, BOLD: p<.00625 corrected for multiple comparisons.

**Figure 4.**
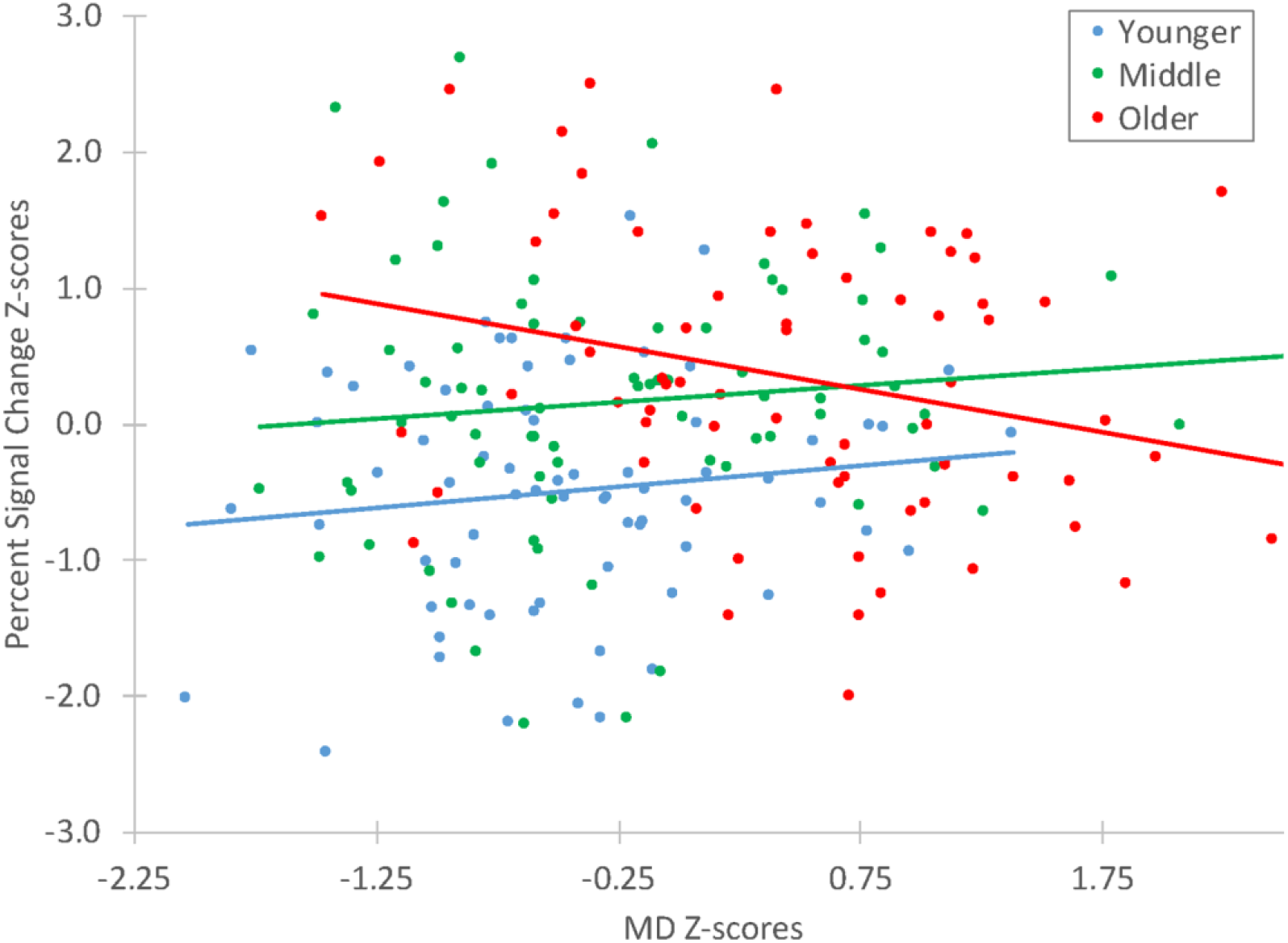
Percent signal change plotted against mean diffusivity (MD) scores for three age groups split by tertiles. Only the oldest age group showed a negative association between the two brain measures, suggesting greater activation is associated with better white matter diffusivity in older adults

## 4. Discussion

This study was the first to examine the brain differences between fluid abilities that decline sharply with age and crystallized abilities that are well maintained with age. The study encompassed twelve fMRI tasks within the same group of participants and that used the fMRI activations to guide white matter tractography for each ability, enabling within-subject comparisons of functional activation and white matter connectivity. Having brain measures for four abilities in the same set of participants was essential for examining the within-subject differences between fluid and crystallized abilities, which eliminated between-subject differences in the comparisons.

### 4.1 Activation patterns

Brain activation patterns associated with each ability were distinct from each other and consistent with previous studies. The two cortical regions uniquely associated with vocabulary matched the regions reported in previous studies that used similar semantic knowledge tasks. Badre et al. (Badre et al., 2005) observed activation in the left inferior frontal and the left middle temporal gyri across a set of four manipulations to select a target with a specified association to the stimulus (“ivy” and “jade” or “ivy” and “league”), similar to the three vocabulary tasks in our study. In contrast, the fluid abilities were distributed across both hemispheres and encompassed large regions of the cortex, all of which were consistent with previous studies. The reasoning-unique regions dominated the anterior and the dorsal lateral prefrontal regions as was reported in an fMRI study on the matrix reasoning task with 0, 1 and 2 relational matrices (Christoff et al., 2001) in which Brodmann areas 9, 10 and 46 showed greater activations in the 2- vs 1-relational contrast. The large parietal activation also resonated with previous studies including a transcranial alternating current stimulation study in which the left parietal cortex was stimulated during performance of the matrix reasoning and the paper folding tasks (Neubauer et al., 2017). The small activation in the right hippocampus supports a previous finding that hippocampal volume correlated with fluid reasoning score in older adults (Reuben et al., 2011). Regions unique to speed were strikingly more subcortical, more inferior and posterior to regions associated with the other abilities, which were expected since the speed ability recruits mainly motoric and visual processes. The orbitofrontal activations also make sense for speed processes as the region has been found in a single-unit recording study in mice to play a role in response conflict resolution (Bryden and Roesch, 2015). The memory ability uniquely recruited fewer regions than both reasoning and speed abilities. The left parahippocampal gyrus and cingulate gyrus were shown to be prominent in a positron emission tomography study using 15O-butanol during performance of the pair associates test (Krause et al., 1999).

Based on the age by ability interactions, z-scores of the percent signal change in functional activation showed that correlations with age for memory, not vocabulary, was significantly different from the other three abilities. A number of functional activation studies have reported increased activations with older age across many cognitive tasks such that activated regions in older adults are typically greater in extent, higher in peak intensity, and consist of additional regions not activated by younger adults (Grady, 2008). With the exception of memory, our results were consistent with these previous studies (Grady, 2008); task-related activation for reasoning, speed and vocabulary followed this age trend in that mean percent signal change showed positive associations with age. The activation for memory, on the other hand, did not correlate with age at all, despite behavioral performance having worsened with age. This was an unexpected result and will be discussed further in the next section.

Task activation associated with the four abilities examined in this study showed that the distinguishing feature for vocabulary from the fluid abilities rests on the qualitative difference in the regional distribution of the brain regions. Vocabulary was associated with the fewest unique brain areas and exhibiting the most left lateralization as expected of an ability that requires extensive language processing.

### 4.2 Memory ability

Even though memory performance scores declined with age and the mean percent signal change for the regions associated with memory correlated positively with memory performance, the brain measures associated with the memory ability did not show all of the expected age trends. MD did show atrophy with older age as expected; for FA, while being negatively correlated with age at p<.05, did not survive multiple comparisons correction, but nevertheless was consistent with existing literature in that white matter tracts decline with age. The more unexpected result was the lack of age trend in the mean percent signal change for the memory ability. Previous studies such as Gron et al. (2003) reported significant positive correlations with age for activations in medial frontal gyrus and negative age correlations in the right pulvinar and precuneus. The discrepancy between these previous studies and our result may be attributed to the set of brain regions captured in our approach, in which only regions unique to each ability were retained, and thus may have extracted only the regions that show consistent activation across age for the memory ability. The unexpected results found for the memory ability warrant further investigation to understand whether the lack of age trends was mainly driven by our analytic approach.

### 4.3 Vocabulary ability

Z-scores for mean FA and MD of the white matter tracts connecting the brain regions associated with each of the four abilities examined in our study followed the expected decline with age, with FA showing negative and MD showing positive correlations with age for each ability. Consistent with the hypothesized direction, correlations between MD and age was the lowest for vocabulary compared to the three fluid abilities, as demonstrated by the highly significant age by ability interaction between vocabulary and each of the three fluid abilities. For FA, despite memory being an anomaly, correlations with age was weaker for vocabulary than for reasoning and speed, with the age by ability interaction being significant at p<.05 but did not exceed the corrected threshold for multiple comparisons. The white matter diffusivity results suggest that white matter connections are better maintained among the regions supporting the processing of vocabulary than at least the reasoning and speed abilities. Given the potential white matter advantage vocabulary has over the fluid abilities, a possible mechanism enabling vocabulary to be better maintained with older age may be that the brain regions supporting vocabulary are more localized and thus do not rely on long distance white matter tracts as much as the fluid abilities do. This specific theory will need to be further investigated in future studies.

### 4.4 Age by MD interaction for Speed percent signal change

Association between percent signal change and MD for speed showed a strong interaction with age group such that percent signal change and MD were negatively correlated only in the oldest age group, suggesting higher percent signal change is associated with more intact white matter tracts, which appears to contradict previous studies relating task activation and DTI measures such as Daselaar et al. (2015). Daselaar et al. (2015) showed that greater retrieval-success activity (hits > misses) was associated with higher FA in older adults for memory and executive function tasks. The apparent contradictory results between the findings may be attributed to two main differences between our studies: 1. different cognitive abilities examined and 2. distinct analytical approaches. Daselaar et al. examined memory and executive function whereas the activation to MD association was found for Speed tasks in our study. Activations for different cognitive processes may show differential relationships with decline in white matter tracts. Furthermore, Daselaar et al. used an event-related design that subtracted activation for misses from hits whereas our study used a block design for Speed tasks, which includes both hits and misses. Thus, these study differences limit the comparability between the results.

### 4.5 Limitations and conclusion

One limitation in the study is the use of only one crystallized ability and that was language based. In order to further understand whether greater localization of brain regions is intrinsic to crystallized abilities, other non-verbal crystallized abilities will need to be contrasted with other fluid abilities within the same participants to fully understand how crystallized abilities are maintained with older age. The lack of associations between most of the brain measures and behavioral performance may be attributed to the myriad of factors contributing to behavioral variability that are not captured by the three brain measures.

In conclusion, our results provided initial evidence supporting the hypothesis that the white matter tracts connecting the brain regions associated with processing of vocabulary are better maintained than that for fluid abilities. Rather than individual differences in processing capacity, as stated in cognitive reserve and resilience theories (Stern, 2012), our results suggest variability in processing capacity within each individual across cognitive abilities, enabled by localized differences in anatomical structures involved across the different abilities.

## Acknowledgements

We are grateful for the support provided by the grants: NIA K01AG051777, NIH/NIA R01 AG026158, and NIH/NIA R01 AG038465.

